# Infectious Reactivation of Cytomegalovirus Explaining Age- and Sex-Specific Patterns of Seroprevalence

**DOI:** 10.1101/102491

**Authors:** Michiel van Boven, Jan van de Kassteele, Marjolein J. Korndewal, Christiaan H. van Dorp, Mirjam Kretzschmar, Fiona van der Klis, Hester E. de Melker, Ann C. Vossen, Debbie van Baarle

## Abstract

Human cytomegalovirus is a herpes virus with poorly understood transmission dynamics. We here provide quantitative estimates of the transmissibility of primary infection, reactivation, and re-infection using age-and sex-specific antibody response data. The data are optimally described by three distributions of antibody measurements, i.e. uninfected, infected, and infected after reactivation/re-infection. Estimates of seroprevalence increase gradually with age, such that at 80 years 73% (95%CrI: 64%-78%) of females and 62% (95%CrI: 55%-68%) of males is infected, while 57% (95%CrI: 47%-67%) of females and 37% (95%CrI: 28%-46%) of males has experienced a reactivation or re-infection episode. Merging the statistical analyses with transmission models, we find that infectious reactivation is key to provide a good fit fit to the data. Estimated reactivation rates increase from low values in children to 2%-6% per year older women. The results advance a hypothesis in which adult-to-adult transmission after infectious reactivation is the main driver of infection.

## Introduction

Human cytomegalovirus (CMV) is a highly prevalent herpesvirus that infects between 30% and 100% of persons in populations throughout the world [1]. Usually thought to be a relatively benign persistent infection, CMV is able to cause serious disease in the immunocompromised and offspring of pregnant women with an active infection [2–5].

CMV also has been implicated in a variety of diseases in healthy persons [4, 6], and may play a role in aging of the immune system [7–10], perhaps thereby reducing the effectiveness of influenza vaccination in older persons [11–13].

Although the importance of CMV to public health is acknowledged, and even though the development and registration of a vaccine has been declared a priority [14, 15], little quantitative information is available on the transmission dynamics of CMV. At present, the only population-level data derive from serological studies, aiming to uncover which part of the population is infected at what age. These studies show that i) a sizable fraction of infants is infected perinatally (before 6 month of age), ii) seroprevalence increases gradually with age and is usually higher in females than in males, and iii) the probability of seropositivity is associated with both ethnicity and socioeconomic status, with non-western ethnicity and lower socioeconomic status being associated with higher rates of seropositivity [1, 16–19].

Person-to-person transmission of CMV to an uninfected person can occur from a primary infected person, or from a person who is experiencing a reactivation episode or has been reinfected [4]. Here, we analyze data from a large-scale serological study to obtain quantitative estimates of the relative importance of these transmission routes [19]. We fit mixture models linked to age- and sex-specific transmission models to the data to study the ability of different hypotheses explaining the serological data. Specifically, we quantify the incidence and transmissibility of primary infection, re-infection, and reactivation. Throughout, our premise is that measurements of antibody concentrations provide information on whether or not a person has been infected, and whether or not re-infection or reactivation have occurred. Persons with low measurements are considered uninfected (susceptible), while persons with intermediate and high antibody concentrations are infected and infected with subsequent re-infection or reactivation, respectively.

We find that approximately 20% of infants is infected at the age of six months in the Netherlands, that seroprevalence (i.e. the fraction of the population that is infected) increases gradually with age to 60% and 70% in 80-year-old males and females, and that the fraction of the infected population with high antibody concentration increases from low values in children to 35% and 55% in elderly male and female persons, respectively. Reactivation rates are found to be substantially higher in females than in males. The analyses show that significant infectious reactivation in adults is necessary to explain the serological data, supporting the notion that infectious reactivation is an important driver of transmission.

The results have implications for control of CMV by vaccination, but also in a broader context, as increased antibody concentrations are an indicator of T cell immune memory inflation, impaired viral control, accelerated immunosenescence, and vascular pathologies (see [10, 20] and references therein).

## Results

### Prevalence estimation

Fig 1 presents the data stratified by sex and age, with fits of the statistical model (Fig 7 for overview). The data and model fit show peaks at low antibody measurements (-2.9 U/ml and ≈-2 U/ml), corresponding to uninfected persons (denoted by S, for susceptible; Methods). In both sexes, there is a third peak at higher measurements (1-3 U/ml) that shifts to higher values with increasing age. This peak is composed of persons who are infected (denoted by L, for latently infected) and persons who are infected with high antibody concentrations (denoted by B, for boosted antibodies). Using the estimated distributions for these classes, we find that classification of persons as uninfected versus infected is near perfect (Youden index: 0.97), while classification of persons with high antibody concentrations is good (Youden index: 0.71; Fig 2, Fig 8). These results correspond well with the threshold for infection of the supplier by the Assay (-0.8 U/ml), and show that our classification with three components is supported by the data (i.e. has high probability of yielding an informed decision).

**Fig 1.**
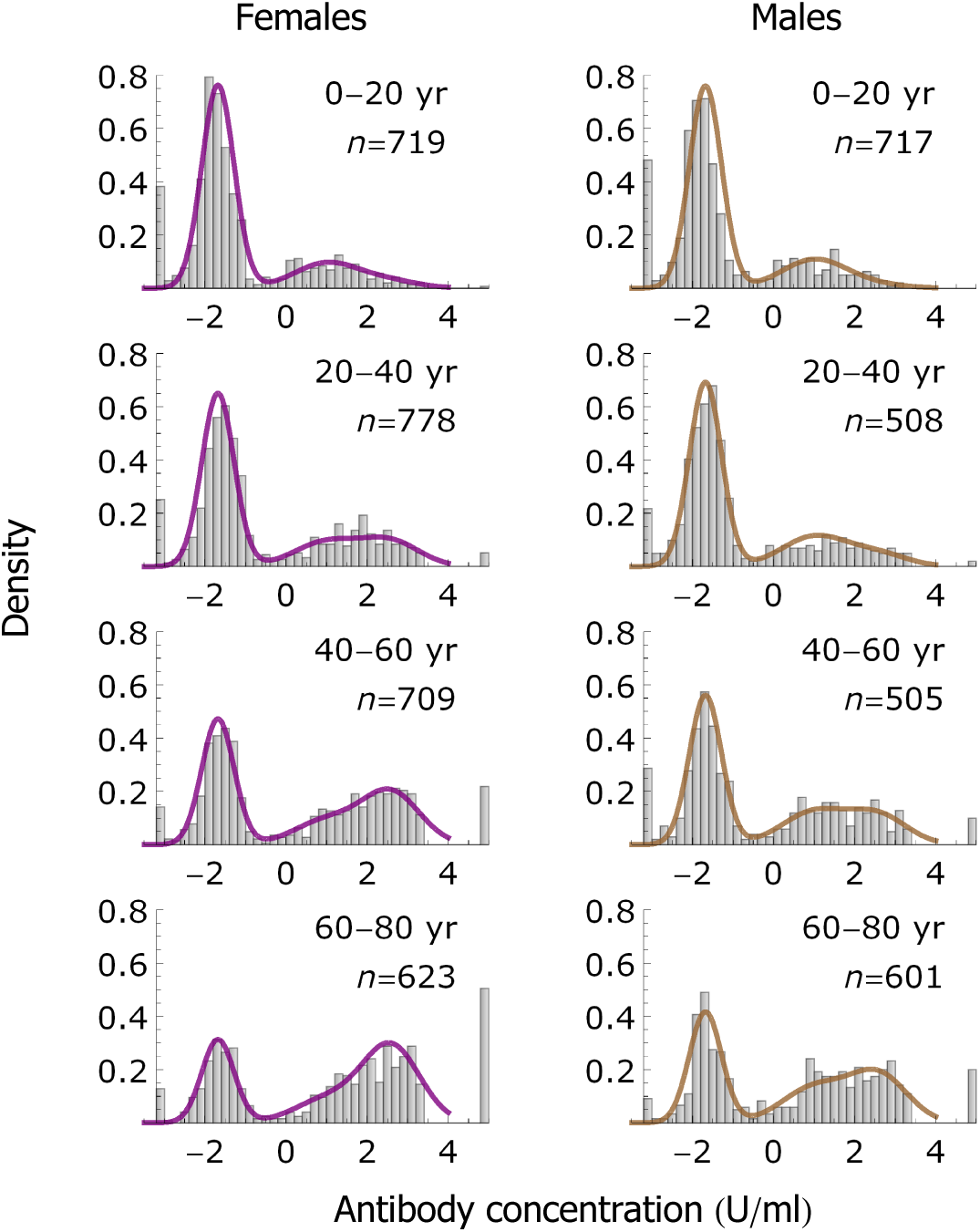
Data and model fit. Data (histograms) and model fit (lines) of antibody concentration measurements by age group and sex. Left- and right-hand panels show results for females (purple) and males (brown), respectively. The leftmost bars at -2.9 contain samples that are assumed uninfected, and the rightmost bars at 4.5 contain samples that are right censored (with concentration >3.41; Methods). Insets show the age group and number of samples.

**Fig 2.**
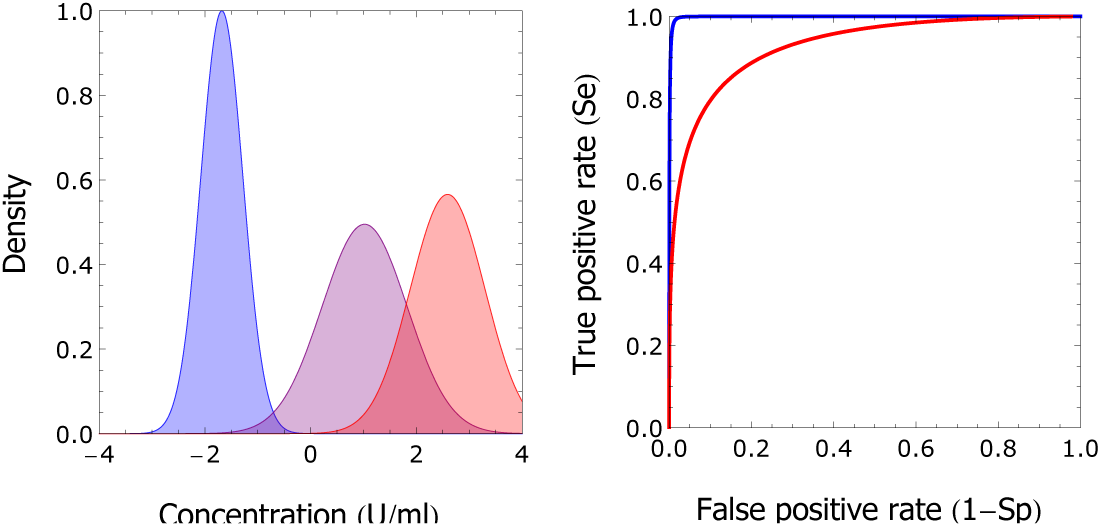
Classification of samples. Shown are the estimated mixture distributions using the parameter medians of the posterior distribution (left-hand panel; blue: susceptible; purple: infected); red: infected with increased antibody concentration), and receiver operating characteristic of binary classifications taking the estimated distributions as ground truth (right-hand panel). Maximal sum of sensitivity and specificity (Se+Sp) for classification of uninfected versus infected persons is 1.97 at antibody concentration -0.70 U/ml, with sensitivity 0.99 and specificity 0.98 (Figure S8). This value corresponds well with the threshold for infection of -0.8 U/ml suggested by the supplier of the assay. Maximal Se+Sp for classification of persons with increased antibody concentration is 1.70 at antibody concentration 1.81 U/ml, with sensitivity 0.84 and specificity 0.87 (Fig 8).

Fig 3 shows the estimated prevalences in females and males as a function of age. The susceptible prevalence decreases gradually with age, from approximately 0.80 in infants (females: 0.81, 95%CrI: 0.77-0.85; males: 0.80, 95%CrI: 0.76-0.84) to 0.27 (95%CrI: 0.22-0.34) and 0.38 (95%CrI: 0.32-0.45) at 80 years in females and males, respectively. In both females and males the latently infected prevalence remains approximately constant, ranging from 0.15 to 0.20 in females and from 0.18 to 0.28 in males. In contrast, the prevalence of persons with increased antibodies increases strongly with age, especially in females. In fact, the prevalence of persons with increased antibodies increases from 0.09 (95%CrI: 0.06-0.13) at 20 years to 0.57 (95%CrI: 0.47-0.67) at 80 years in females, and from 0.04 (95%CrI: 0.03-0.07) to 0.37 (95%CrI: 0.28-0.46) in males. Hence, in older persons the prevalence of persons with increased antibodies is 54% (or 20 per cent points) higher in females than in males.

**Fig 3.**
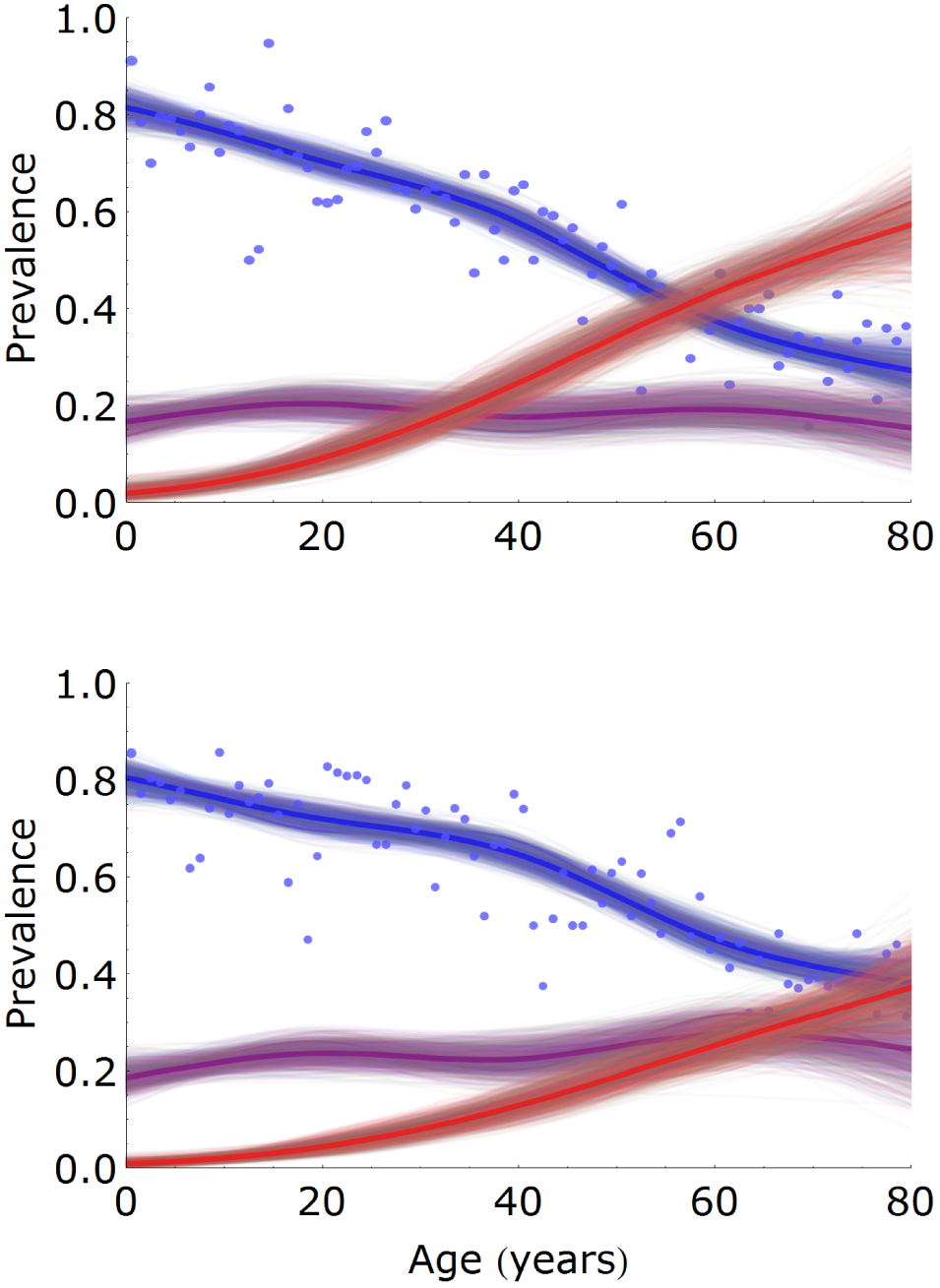
Estimation of age- and sex-specific prevalence. Prevalence estimates are presented for females (top) and males (bottom), and for classes of low (susceptible, blue), medium (latently infected, purple), and high (latently infected with increased antibodies, red) antibody measurements. Shown are 1,000 samples from the posterior distribution (thin lines) with posterior medians (bold lines). Dots indicate the fraction of samples in each year-class that would be classified as uninfected with the cut-off specified by the supplier of the assay. The number of samples per year-class is approximately 35 and 30 for females and males, respectively.

Of particular interest is the prevalence of infection in females of childbearing age, as this group is at risk of transmission to the fetus or newborn. Using the above analyses, we find that the prevalence of infection (i.e. the combined prevalence in the L and B classes) is 0.30 (95%CrI: 0.27-0.33) in 20-year-old females and 0.42 (95%CrI: 0.39-0.46) in 40-year-old females. If we combine these figures with the observation that approximately 20% of children infected at six months of age, and that less than 5% of children in the Netherlands in 2007 had a mother under 20 years or over 40 years, we deduce that the probability of perinatal transmission could be between 0.20/0.42=0.48 and 0.20/0.30=0.67, with the exact figure depending on the distribution of ages at which mothers give birth. In addition, one could envisage that the highest risk of (severe) infection of the fetus or newborn is when mothers are infected or experience a reactivation episode. The estimated rates at which susceptible females of 20 and 40 years are infected are 0.0055 per year (95%CrI: 0.0036-0.0077) and 0.0092 per year (95%CrI: 0.0069-0.011) per year, respectively. The rates at which latently infected females of 20 and 40 years are re-infected or experience a reactivation episode are of similar magnitude, and are estimated at 0.0059 per year (95%CrI: 0.0038-0.0086) and 0.0093 per year (95%CrI: 0.0064-0.012), respectively. The overall rates of infection, reactivation, and re-infection in 20 and 40 year-old females are given by the sum of the above estimates, and are approximately 1% and 2% per year, respectively.

### Estimation of reactivation and re-infection rates

Naive estimates of the reactivation and re-infection rates can be obtained by transformation of the spline prevalence estimates (Methods), indicating that infectious reactivation is required to explain the serological data (Fig 9). Estimation with the transmission model confirms these findings, and shows that a model without reactivation fits the data worse (ΔAIC=426.3) than either a model with reactivation and re-infection (full model), or a model with reactivation but no re-infection (Table 1; Methods). A fourth model with reactivation and re-infection that is not infectious (*β*_2_=0) performs better than the model without reactivation, but worse than either the full model or the model with no re-infection (ΔAIC=75.2).

**Table 1.** Model selection of transmission scenarios. For each of four model scenarios the number of parameters, maximal log-likelihood ℓ_max_, and AIC difference with the best fitting model are given.

| Scenario | $n$ | $\ell_{max}$ | $\Delta AIC$ |
| --- | --- | --- | --- |
| Full model | 9 | -7195.2 | 2.0 |
| No re-infection | 8 | -7195.2 | 0 |
| No reactivation | 3 | -7413.3 | 426.3 |
| Reactivation/re-infection not infectious | 8 | -7233.8 | 75.2 |

The prevalences estimated with the mixture model and the corresponding maximum likelihood estimates of the transmission models are given in Fig 10, and show that a transmission model without reactivation overestimates the prevalence of infection, especially in children. Overall, models without infectious reactivation have low empirical support, while there is no decisive statistical evidence in favor of either a model with infectious reactivation and infectious re-infection (ΔAIC= 2:0) or a model with infectious reactivation only (highest AIC; [21]). These results are supplemented and supported in a Bayesian analysis of the latter two models (Methods). Using the Watanabe Akaike information criterion (WAIC) that better gauges out-of-sample predictive performance than AIC we find similar performance of both models (WAIC=14,410.4 for the full model versus WAIC=14,409.1 for the model without re-infection; [22, 23]).

Inspection of the parameter posterior medians of the two best-performing models shows that both yield similar estimates of the transmissibility of primary infection (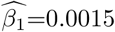 per year (95%CrI: 6.2.10^−5^-0.0072) for the full model and 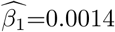 per year (95%CrI: 4.9.10^−5^-0.0071) for the model without re-infection), and nearly identical estimates of the transmissibility of reactivation and re-infection (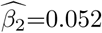
 per year (95%CrI: 0.045-0.057) in both scenarios). In the model without re-infection the proportionality parameter governing the re-infection rate (the probability of re-infection in a contact that would lead to infection if the contacted person were uninfected) is zero by definition, and in the full model the proportionality parameter is dominated by the prior distribution *U*(0, 1), indicating that it cannot be estimated with meaningful precision from the data (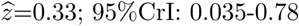).

Estimates of the reactivation rates are quantitatively close in models with reactivation (Fig 4). Reactivation rates generally increase with increasing age, and are substantially higher in females than in males. In the full model, the estimated reactivation rate is 0.015 per year (95%CrI: 0.0064-0.025) in 0-20 year-old females, which doubles to 0.030 per year (95%CrI: 0.019-0.041) in 20-50 year-old females, and then increases further to 0.042 per year (95%CrI: 0.021-0.062) in 50+-year-old females. The corresponding reactivation rates in males are 0.0022 per year (95%CrI: 0.0010-0.0074), 0.019 per year (95%CrI: 0.0098-0.029), and 0.012 per year (95%CrI: 0.0015-0.023). In the model without re-infection these estimates are 0.019 per year (95%CrI: 0.012-0.028), 0.033 per year (95%CrI: 0.024-0.043), and 0.045 per year (95%CrI: 0.026-0.064) in females, and 0.0052 per year (95%CrI: 0.0019-0.010), 0.021 per year (95%CrI: 0.014-0.030), and 0.015 per year (95%CrI: 0.0016-0.027) in males. Hence, estimates of reactivation rates are higher and slightly more precise in the model without re-infection than in the model with re-infection.

**Fig 4.**
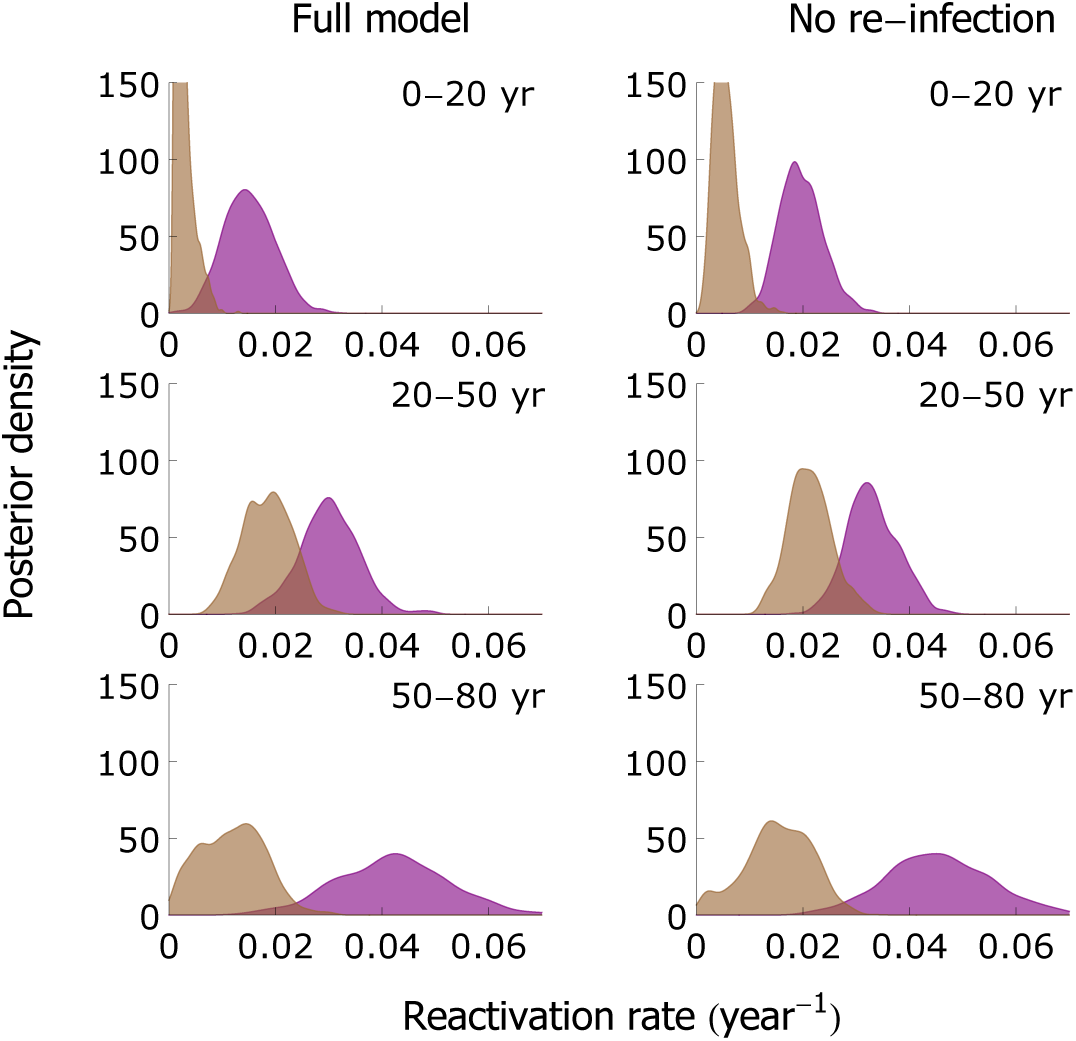
Estimation of age-specific reactivation rates. Shown are kernel-smoothed estimates of the reactivation rates in persons aged 0-20 years (top), 20-50 years (middle), and 50-80 years (bottom) in the full model (left-hand panels) and model without re-infection (right-hand panels; Table 1). Estimates for females and males are shown in purple and brown, respectively.

In both models with reactivation, estimates of the force of infection increase from approximately 0.01 per year in the youngest age group to 0.012-0.014 per year in 10-15 year-old girls (Fig 5). Owing to the slightly higher contact rates in females than in men, the estimated force of infection is usually a couple percentage points higher in females than in males in the age groups 10-25 years [24]. In older age groups, estimates of the forces of infection decrease to lower values (0.005-0.01 per year). Noteworthy, the extreme age-specific differences in the force of infection usually observed for directly transmitted infectious diseases, with high infection rates in children and much lower rates in adults, are much less pronounced here due to infectious reactivation combined with age-assortative mixing (Fig 10) [24–26].

**Fig 5.**
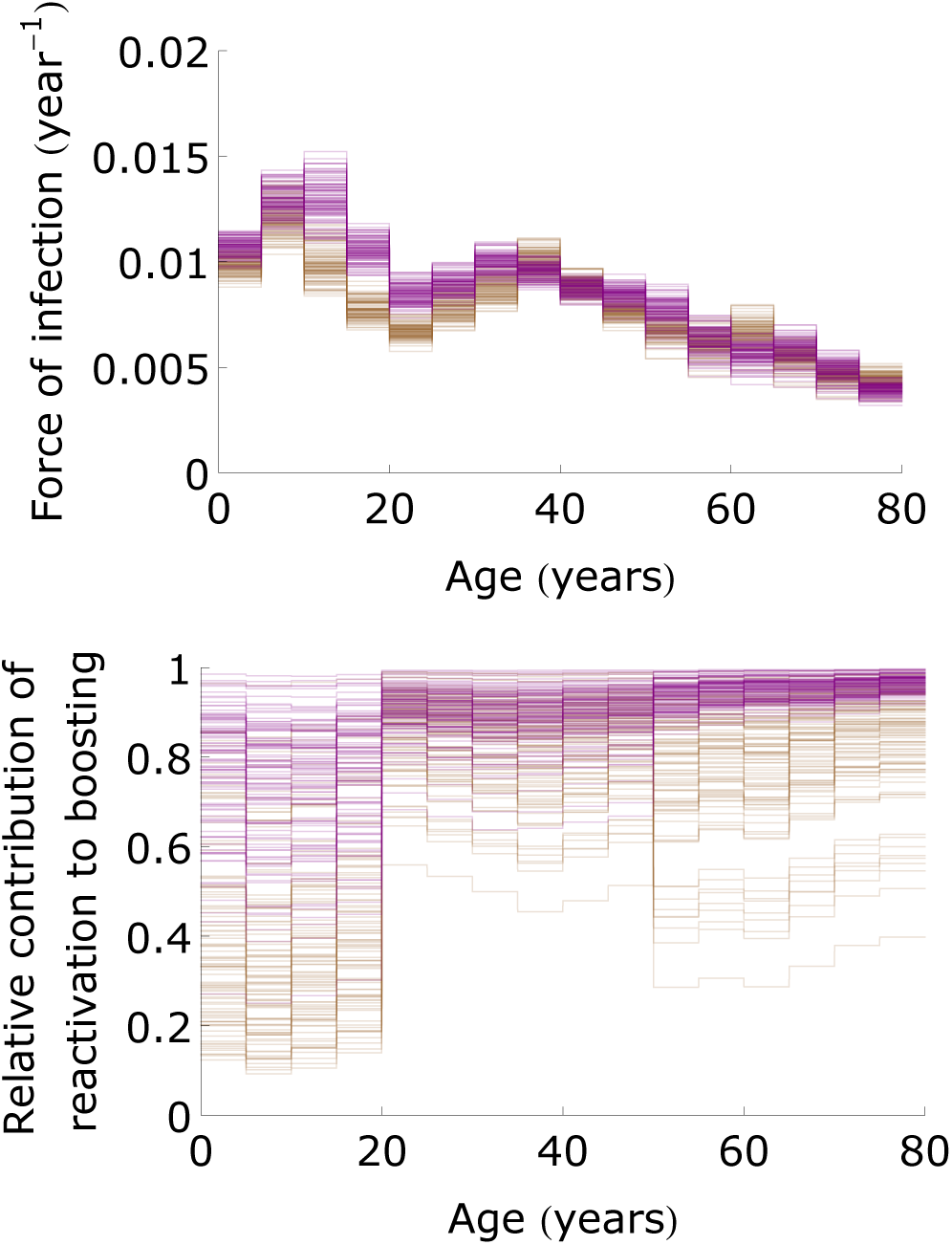
The force of infection and contribution of reactivation to antibody boosting. Shown are estimates of the forces of infection in females and males (top; purple: females; brown: males) and relative contributions of reactivation to antibody boosting 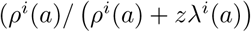 with 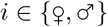) (bottom) in the full model as a function of age. Both panels show 100 samples from the posterior distribution.

Further, estimates of re-infection rate (*z*λ(a)) are considerably smaller than estimates of the reactivation rates 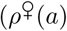 and 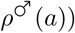 because the estimated force of infection (λ(a)) is small in all age groups (Fig 5). Hence, re-infection contributes little to boosting of the antibody concentrations in those age groups where most of the boosting occurs (>20 years; Fig 3).

## Discussion

Our study of population-wide serological data shows that IgG antibody concentrations contain a wealth of information on the transmission dynamics of CMV. Specifically, the analyses reveal that (i) the prevalence of CMV increases gradually with age such that at old age the majority of persons in the Netherlands are infected; (ii) except for the very young, prevalence of CMV is systematically higher in females than in males. This is mainly due to a higher incidence of infection in adult women than in adult men of similar age; (iii) antibody concentrations in seropositive (i.e. infected) persons increase monotonically with age, especially in women; (iv) the above findings (i)–(iii) cannot be explained by simple transmission models in which only primary infection is infectious. This is caused by the fact that transmissibility of primary infection determines the rate at which age-specific prevalence increases; if transmissibility of primary infection would be high then a high prevalence of infection is expected in children (Fig 10). In other words, the fact that seroprevalence increases gradually with age puts an upper bound on the force of infection, and this in turn constrains the transmissibility of primary infection to low values.

While aforementioned findings (i)–(iii) have been noticed before in other settings ( [1] and references therein, [19]), our analyses are the first to provide precise estimates using a large population sample. Moreover, the analyses advance a transmission hypothesis based on the notion that reactivation contributes significantly to the transmission dynamics of CMV. Since prevalence of infection has been shown to increase in a gradual manner in several other studies [1], this explanation may not be restricted to the Dutch situation, but hold in general. Underpinning this hypothesis, next to the well-known observations of shedding of CMV in breast milk and cervical material in the third trimester of pregnancy [27–29], detectable virus also has been found in some healthy persons under 70 years and in all persons over 70 years in one study [31], while in another study CMV DNA has been detected in urine of the majority of older persons [32]. These findings add to our belief that infectious reactivation may be a plausible explanation for the patterns of infection observed in serological data.

The main implication is that the majority of CMV infections may not be caused by transmission among children after primary infection, even though levels of shedding can be high in infants [28, 33], but rather by older persons who go through a reactivation episode. As a result, infectious reactivation is expected to be an important driver of CMV transmission in the population. This contrasts with common childhood diseases such as measles, mumps, rubella, and pertussis, for which infection in unvaccinated populations generally occurs at a young age and children are the drivers of transmission. It also contrasts with other herpes viruses such as varicella zoster virus and Epstein-Bar virus for which well over 50% of the population is infected at the age of 10 years [25]. It may be comparable with other herpes viruses such as HSV1 and HSV2, which show a slowly increasing age-specific seroprevalence [34]. A corollary is that primary CMV infection alone is unlikely to be able to maintain sustained transmission in the population. In fact, further analyses based on the current parameter estimates suggest that primary infection coupled with infectious reactivation would not be sufficient to ensure persistence, as the estimated basic reproduction number is smaller than the threshold value 1 (Methods; Fig 12). This, in turn, indicates that perinatal transmission is required for persistence.

With infectious reactivation and perinatal infection being putative drivers of transmission, it is to be expected that elimination by vaccination may prove more difficult than for directly transmitted pathogens, as it will require the pool of latently infected persons to dwindle to zero by demographic turnover. This can take up to the lifetime of one generation, and perhaps more if vaccination cannot prevent perinatal transmission to infants who are too young for vaccination. Thus, a question is whether vaccination formulations and strategies exist that minimize the probability of transmission to young infants. This is all the more of importance as a main source of morbidity is by congenital infection, and the timescale on which reductions in congenital disease are expected determines the projected health impact of vaccination [35]. In this context, next to the ability of a vaccine to prevent infection it may perhaps be equally important that a vaccine is able to reduce the probability of reactivation. Such reductions are likely mediated by T-cell responses of the host, and several (but not all) vaccines under development are expected to induce boosting of T-cell immune responses [36–38].

A number of limitations and assumptions deserve scrutiny. First, the transmission model analyses assume that the population is in endemic equilibrium. For a single cross-sectional data set such as considered in the present study this assumption is unavoidable if one does not want to introduce additional parameters that cannot be estimated by the data. Reassuringly, the patterns of infection present in the serological data have been found in several serological studies carried out in high-income countries over the past decades [1]. Also, no systematic patterns of increasing or decreasing seroprevalence over time have been found, and this is further reason to believe that there have not been major changes in the epidemiology of CMV over time [1]. Second, we assume that antibody measurements not only give information on CMV infection status, but also whether or not reactivation/re-infection have taken place.

Unfortunately, there is no direct empirical evidence available confirming or falsifying this assumption, and this is an area where in-depth comparison of the infection and immune status of persons with low and high antibody concentrations is urgently needed. Third, the analyses assume that person-to-person transmission is proportional to observed human contact patterns (Fig 11; [24, 39]). Although this assumption is commonly made and has met with considerable success (e.g., [39–42]), it is conceivable that transmission of CMV does not abide by the social contact hypothesis, and that a more complex contact structure would be able to explain the patterns of seroprevalence in a simple transmission model. To investigate the impact of the contact structure, we have re-analyzed all transmission model with a uniform contact structure, and found that a model with infectious reactivation provides the best fit to the data (not shown). By extension, it also remains a possibility that the patterns of infection are generated by a complex pattern of susceptibility increasing with age. Again, evidence for or against this possibility is lacking.

Throughout, the analyses are based on the observation that, at the population level, CMV IgG antibody concentrations are well-described by three distributions. We have exploited this observation to build a transmission model with three classes, pertaining to persons who are uninfected, infected, or infected after reactivation or re-infection. In this model, a person experiences at most one reactivation or re-infection event during its lifetime. In reality, it is more likely that such reactivation and/or re-infection events may occur more often. Unfortunately, it is not possible to statistically identify the parameters in models that would include multiple reactivations/re-infection events. As a result, transitions from the infected class to the infected class with increased antibodies may in fact be the result of multiple underlying reactivation or re-infection events, and the infectiousness parameter of reactivation and re-infection should be interpreted as a compound parameter describing the overall impact of multiple reactivations and re-infections. This may help explain why estimated infectiousness of primary infection is much lower than estimated infectiousness after reactivation or re-infection, even though it is known that prolonged and high-level virus shedding can occur in bodily fluids after primary infection in children [28, 29].

## Materials and Methods

### Study design

The analyses make use of sera from the PIENTER2 project, a cross-sectional population-based study carried out in the Netherlands in 2006-2007. Details have been published elsewhere [19]. Briefly, 40 municipalities distributed over five geographic regions of the Netherlands were randomly selected with probabilities proportional to their population size, and an age-stratified sample was drawn from the population register. A total of 19,781 persons were invited to complete a questionnaire and donate a blood sample. Serum samples and questionnaires were obtained from 6,382 participants. To exclude the interference of maternal antibodies, we restrict analyses to sera from persons older than 6 months (6,215 samples). We further select Dutch persons and migrants of Western ethnicity to preclude confounding by ethnicity (5,179 samples) and stratify the data by sex [19], yielding 2,842 and 2,337 samples from female and male participants, respectively. The data are available at GitHub (github.com/mvboven/cmv-serology).

### Ethics

The study was approved by the Medical Ethics Testing Committee of the foundation of therapeutic evaluation of medicines (METC-STEG) in Almere, the Netherlands (clinical trial number: ISRCTN 20164309). All participants or their legal representatives had given written informed consent. Details of the study are given elsewhere [30].

### Antibody assay

We use the ETI-CYTOK-G PLUS (DiaSorin, Saluggia, Italy) Elisa to detect CMV-specific IgG antibodies. The assay yields continuous measurements (henceforth called’antibody concentration’) and has an upper limit of 10 units/ml in our test setting. A small number of samples is right-censored at this limit (140 persons). According to the provider of the assay, samples with measurement lower than 0.4 units/ml should be classified as uninfected, while samples with measurement ≥0.4 units/ml should be classified as infected.

### Mixture model

The data are analyzed using a mixture model with sex- and age-specific mixing functions. A Box-Cox transformation with parameter 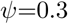 yields a distribution of antibody measurements of samples with low antibody concentration that is approximately Normal, and we henceforth base all analyses on the transformed measurements (denoted by U/ml). The estimated prevalences are robust to this transformation, which merely serves to ensure that the data are well-described by Normal mixture distributions (not shown). We distinguish three distributions, describing samples with low (susceptible, S), intermediate (latently infected, L), and high (increased antibodies, B) antibody concentrations. The L and B distributions are modeled by a Normal distribution with means and standard deviations independent of age and sex, and the S distribution is modeled by a mixture of a spike and a normal distribution (see below).

Right-censoring is applied to the 140 samples above the upper limit of 3.41 U/ml (10 units/ml on the original scale). As there appears a spike at -2.91 U/ml (0.0001 units/ml on the original scale) in the data (263 persons), we model the S component by a mixture of a spike at this value and a Normal distribution (i.e. an inflated normal distribution). In this way, samples with concentration at the spike belong to the susceptible component with probability 1.

We model the probability of each of the three outcomes in terms of log-odds, taking the probability of being in the S component as reference. This allows us to write the log-odds of being in component L or B as linear functions of age and sex. The design matrix of the resulting multinomial logistic model consists of natural cubic splines with interior knots at 20, 40 and 60 years and boundary knots at 0 and 80 years. Hence, the mixing functions (prevalences) have flexible shape, which allows these to be optimally informed by the data. In the results, sex is put in the model as main effect, as analyses show no noticeable improvement in fit when including age by sex interaction (not shown).

We estimate parameters in a Bayesian framework using R and JAGS [43, 44]. Non-informative Normal prior distributions are set on the means of the three component distributions (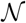 (0, 0.001)). Label switching is prevented by prior ordering of the means. The precisions of the components are given flat Gamma prior distributions (Γ(0.5, 0.005)). The spline parameters are also given non-informative Normal prior distributions (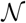 (0, 0.001)). We apply a QR-decomposition to the design matrix to improve mixing and run 10 MCMC chains in parallel, yielding a total of 10,000 samples. Although mixing of the unthinned samples is already satisfactory, we apply an additional 1/10 thinning, giving a total of 1,000 samples from the posterior distribution.

### Classification

The distributions characterizing the subpopulations with low (susceptible, S), intermediate (latently infected, L), and high (increased antibodies, B) antibody concentrations allow assignment of probabilities to individual samples. For any single observed sample we can calculate the probability that the corresponding person is susceptible, latently infected, or latently infected with increased antibodies. In detection theory, the specificity (the probability of correctly classifying a negative subject) and sensitivity (the probability of correctly classifying a positive subject) are used to characterize the power of a detection procedure. The relation between sensitivity and specificity may be graphed with antibody concentration specifying a cut-off for binary classification as a parameter in a receiver operating characteristic (ROC) graph [42, 45, 46]. Here, we investigate the ability of the mixture analyses to distinguish uninfected (S) from infected (L+B) samples, and to distinguish samples with increased antibody concentration (B) from uninfected and latently infected samples (S+L). The results are laid down in Fig 2 and Fig 8.

### Transmission model

Next to the statistical analysis using mixture models, we analyze the data with transmission models. The transmission models take the estimated mixture distributions or, more precisely, the medians of the posterior distribution as input. In line with the above, we focus on a sex- and age-structured model in which persons are probabilistically classified in one of three classes, viz. uninfected (S), latently infected (L), and latently infected after reactivation or re-infection (B). As the infectious period is short relative to the lifespan of the host (weeks versus decades) we do not explicitly model the infectious periods, and assume that the transitions from the S to the L and from the L to the B classes occur instantaneous (the short-disease approximation, [47]). Further, we focus on the endemic equilibrium of the transmission model so that all variables are time-independent [47, 48]. Fig 6 shows a schematic of the model. For sexes 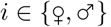, the age-dependent differential equations are given by

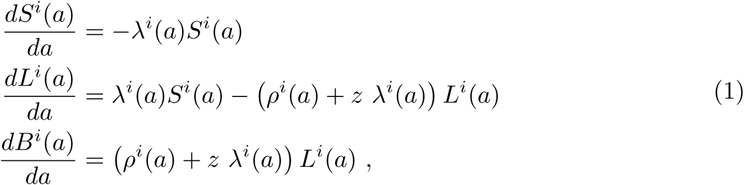

with forces of infection

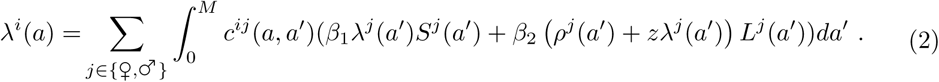

In Eqs (1)-(2), *zλ^j^*(*a*) and Eq *ρ^j^*(*a*) are the age-specific re-infection and reactivation rates, *z* is the susceptibility to re-infection of latently infected persons relative to the susceptibility of uninfected persons 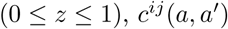 represents the contact rate between persons of age *a′* and sex *j*, and those of age *a* and sex *i* [24, 39], *β_1_* and *β_2_* are proportionality parameters determining the transmissibility of primary infection and reactivation/re-infection, and *M* is the maximum age. As the data do not extend beyond 80 years we take *M* = 80 years. Notice that 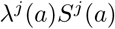 and 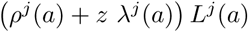 are the incidence of primary infection and the incidence of reactivation and re-infection, so that 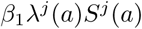 and 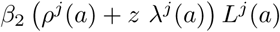 are the infectious output generated by primary infection and reactivation/re-infection, respectively [47].

As in earlier studies, contact rates (contact intensities in the notation of [24] are hard-wired into the model using data on social contact patterns, thereby adopting the social contact hypothesis [24, 25, 39]. Here we use the mixing matrix of an analysis of the data from the Netherlands gathered in 2006/2007 with demographic composition of the Dutch population in 2007 [24]. See Fig 11 and GitHub (github.com/mvboven/cmv-serology) for the contact matrix and demographic data.

The differential equations can be solved in terms of the forces of infection using the variation of constants method. Here we assume, based on results of the mixture model, that a non-negligible fraction of infants is infected in the first six months of life and the fraction infected is equal in female and male infants [19]. Hence, we have 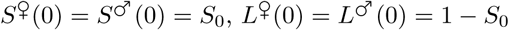 and 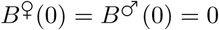 as initial conditions, and the solution of (1) is given by

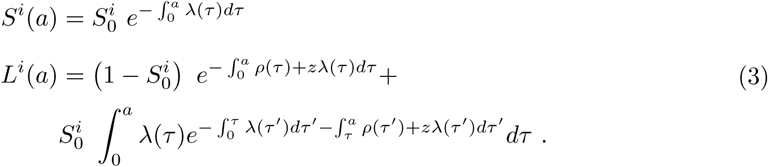

**Fig 6.**
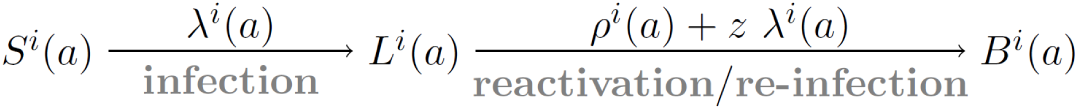
Schematic of the model. *S^i^*(*a*) denotes the age-specific proportion of uninfected persons of sex 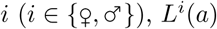 represents the age-specific proportion of latently infected persons, and *B^i^*(*a*) is the age-specific proportion of infected persons with increased antibody concentration. The infection and re-infection rates are given by 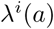 and 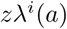, and the reactivation rates are given by *ρ^i^*(*a*).

Insertion of Eq (3) in Eq (2) yields two integral equations for the age-specific forces of infection in females and males [25, 26, 49, 50]. These equations cannot be solved explicitly in general. It is possible, however, to solve the equations for specific functions. Here we assume that transition rates and contact rates are constant on predefined age-intervals, and insert the explicit solution of Eq (3) in terms of force of infection in Equation (2) (see below).

### Naive estimation

Before turning attention to estimation based on the full transmission model, we would like to remark that naive estimates of the force of infection and reactivation rate that do not take the infection feedback (i.e. Eq 2) into account can be obtained by transformation of the spline prevalence estimates. Specifically, rearranging the model equations (Eq (1)) yields the following expressions for the force of infection and reactivation rate

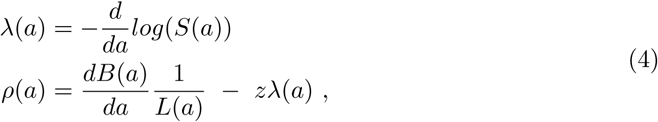

where superscripts denoting sex have been dropped for notational convenience. Since the prevalences *S*(*a*), *L*(*a*), and *B*(*a*) are known (the spline estimates) and can be differentiated, this immediately yields an estimate of the force of infection *λ*(*a*) and subsequently also an estimate of the reactivation rate *ρ*(*a*) (if *z* is assumed known). Notice that these estimates are not necessarily compatible with the transmission model. In particular, the rates so estimated are not necessarily positive (Fig 9).

Assuming that re-infection is negligible (z=0), we find that estimates of the forces of infection are low up to the age of 40 years in both females and males (generally <0.01 per year), increase to higher values in adults aged 40-60 years (0.01-0.03 per year), and are variable in older adults (Fig 9). Estimates of the reactivation rates increase from low values in children to 0.03-0.08 per year in 40-year-old females, and to approximately 0.02 per year in 40-year-old males. At older ages, estimates of the reactivation rates are variable, and have the tendency to drop to slightly lower values. If we take the alternative extreme that primary infection and re-infection occur at identical rates (*z* = 1), estimates of the reactivation rates are lower but the general pattern of substantial reactivation in adults remains (Fig 9).

### Numerical solution of the forces of infection

For statistical analysis based on the full transmission model (Eqs 1)-(2) we assume that reactivation and contact rates are constant in certain predefined age-intervals. From Eq (2), it then follows that the force of infection is piecewise constant as well. Throughout, we consider age intervals of fixed size Δ*a* = 5 years, so that the limits of the 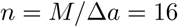 age classes are defined by the vector 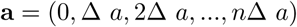. Hence, the *j*-th class (*j* = 1,…, *n*) contains all persons with age in the interval 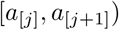, where *a*_[*j*]_ denotes the *j*-th element of a. Subsequently, the forces of infection λ^*i*^(*a*) and reactivation rates *ρ^i^*(*a*) are replaced by their counterparts 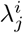 and 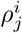. Similarly, 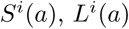 and *B*^*i*^(*a*) are replaced by 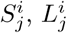 and *B_j_^i^*. Insertion in Eq (3) and integrating over the (constant) rates yields

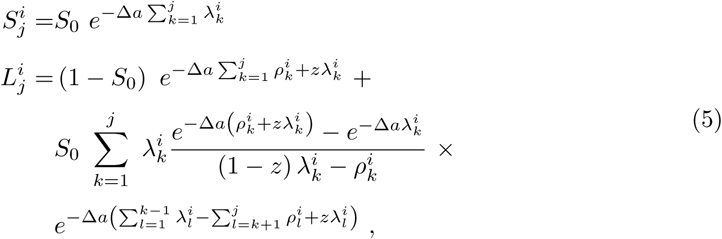

where 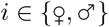 and 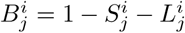. Insertion of Eq (5) in Eq (2) and making use of the fact that the cumulative incidences of infection and reactivation/re-infection in age class *j* are given by 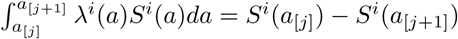 and 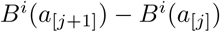, yields 2*n* = 32 equations (*n* = 16 per sex) for the forces of infection. These equations can be solved numerically. Here we use a Quasi-Newton (secant) method to solve the equations.

### Likelihood and estimation

With transition rates and forces of infection at hand, we calculate the log-likelihood of the data. Here, the log-likelihood of each observation is given by a mixture distribution. For instance, the log-likelihood contribution of a sample with antibody measurement *c* in a person of sex *i* and age *a* is given by log 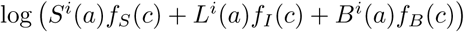, where *S^i^*(*a*), *L^i^*(*a*), and *B^i^*(*a*) are the age specific prevalences in sex *i*, and *f_S_*(*c*), *f_L_*(*c*), and *f_B_*(*c*) are the densities of the mixture distributions at antibody concentration *c*.

To reduce computation times and enable better comparison of the prevalence estimates of the statistical and mechanistic models, we take the posterior medians of the component distributions of the logistic model as inputs in the transmission model. Hence, the transmission model takes the component distributions as given and provides estimate the prevalences via the transmission and reactivation rates. Here, based on preliminary analyses, reactivation rates are modeled by piecewise constant functions with steps at 20 and 50 years, i.e. with constant rates on the intervals [0, 20), [20, 50), and [50, 80) years. Hence, the reactivation rates are characterized by three parameters in each sex, viz. 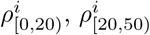 and 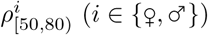.

We employ a Markov chain Monte Carlo (MCMC) method to obtain maximum likelihood (ML) estimates in a pre-screening of models as well as Bayesian estimates for a subset of models that perform well in the pre-screening. For this purpose, we take non-informative (improper) uniform prior distributions for all parameters. Steps in the estimation procedure are as follows. First, for new parameters 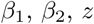 or 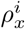 we solve the 2*n* = 32 equations for the forces of infection 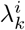. Second, we use the forces of infection to calculate the age-specific prevalences *S^i^*(*a*), *L^i^*(*a*), and *B^i^*(*a*). Third, the age-specific prevalences are used to calculate the log-likelihood. Fourth, the new parameters are accepted or rejected, and the above steps are repeated. Throughout, updating of parameters is based on a single component random-walk Metropolis algorithm using Normal proposal distributions with the current value as mean and standard deviations tuned to achieve acceptance ratios in the 20%-50% range. In the Bayesian analyses, output is generated for 5,000 cycles after a burn-in of 1,000, and a thinned sample of 1,000 is used for analysis. Convergence of chains is assessed visually. Parameter estimates are represented by posterior medians, and bounds of 95% credible intervals are given by 2.5 and 97,5 percentiles. For model selection, we report the Watanabe Akaike Information Criterion (WAIC) (using p_WAIC2_, see [22, 23]). We haved performed a pre-screening of models using ML since the Bayesian analysis does not always yield properly converged chains for poorly fitting models. Here, we use a simulated annealing method in which the Metropolis acceptance probability *p* is replaced by *p^1/T^*, where 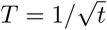 is the temperature at iteration *t*. In this manner, proper ML estimates can be obtained within hours (∼500 iterations). Comparison of models in the pre-screening is based on the Akaike Information Criterion (AIC). The analyses are performed with Mathematica 10.0.

In the main text we consider a suite of simplifications of the full model specified by Equations (1)–(2). In the simplification we assume that (i) there is no re-infection (*z* = 0), (ii) there is no reactivation 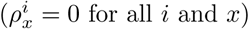, or (iii) reactivation and re-infection are not infectious (*β*_2_ = 0).

### Reproduction numbers

With estimates of key parameters at hand we can further our understanding of the transmission dynamics of CMV, e.g., by calculation of the basic and type reproduction numbers [47]. These reproduction number give insight in the drivers of transmission, and also in the control effort required for elimination or eradication. Full calculation is not straightforward, as CMV can be transmitted after primary infection, after reactivation, and perinatally from mother to offspring. While our analyses have yielded estimates of the former two, formal estimates of perinatal transmission are still lacking. Furthermore, calculation of reproduction numbers implicitly assume a stable host demography and known female fertility distribution function [51]. To obtain partial insight of the impact of the transmission by sex, age and transmission route, we assume type 1 demography (everybody lives exactly to the age of *M* = 80 years), and calculate reproduction numbers in the presence of direct transmission and transmission after reactivation but without perinatal transmission. Central in the calculations are the so-called kernels 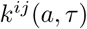 of the next-generation operator, which determine the number of cases of sex *i* and age *a* generated by an infected person of sex *j* and age *τ*. In our case, a straightforward calculation [47] shows that 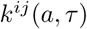 is given by

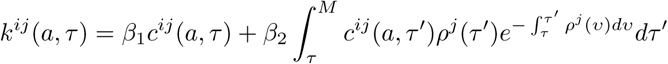

Discretizing the above kernel while making the simplifying assumption that only a single event can happen in any one age group, we obtain approximate estimates of the sex- and age-specific reproduction numbers, collected in the so-called next-generation matrix (Fig 12). Subsequently, we derive estimates of the basic reproduction number *R_0_* and the reproduction numbers between the sexes *R^ij^* as the dominant eigenvalues of the next-generation matrix and the sex-specific next-generation matrices, respectively. These calculations give the following results: *R_0_* is estimated (by its posterior median) at 0.83 (95%CrI: 0.78-0.93), and the type reproduction numbers are estimated at 0.57 (95%CrI: 0.53-0.62) for transmission from female to female, 0.45 (95%CrI: 0.42-0.50) for transmission from female to male, 0.30 (95%CrI: 0.25-0.36) for transmission from male to female, and 0.31 (95%CrI: 0.28-0.40) for transmission from male to male. Hence, the above estimates indicate that females contribute more to overall transmission than males because of the higher estimated rates of reactivation in females than in males, while CMV would be unable to persist in the population in the absence of perinatal transmission (as *R_0_* < 1).

Fig 12 further shows that males older than 50 years do not infect many persons because they have a low reactivation rates and low contact intensities. Infected females, especially young females, are expected to infect significantly more persons, the reason being that their reactivation rates and contact intensities are higher. The observation that both males and females generally infect persons that are older than themselves can be explained by the fact that reactivation generally occurs years after primary infection, and that such reactivation is expected to produce infection in persons who have similar age at the moment of reactivation, due to highly age-assortative mixing patterns (Fig 11).

The above analyses can be generalized and made more precise by using full vital statistics of the population (i.e. by using historical age-specific birth rates and anticipated future trends, and historical and anticipated age- and sex-specific mortality rates), but this is beyond the scope of the current study. The above partial analyses provided here do provide evidence that CMV needs both perinatal transmission and transmission after reactivation to be able to persist in the population.

## Acknowledgments

We thank Can Keşmir for discussion, and the persons included in the PIENTER2 study for their participation.

## Competing interests

This work was supported by the Dutch Ministry of Health, Welfare and Sport and the Netherlands Organisation for Scientific Research (grants 645.000.002 and 823.02.014). The funders had no role in study design, data collection and analysis, decision to publish, or preparation of the manuscript. All authors declare that no competing interests exist.

## Supplementary Figures

**Fig 7.**
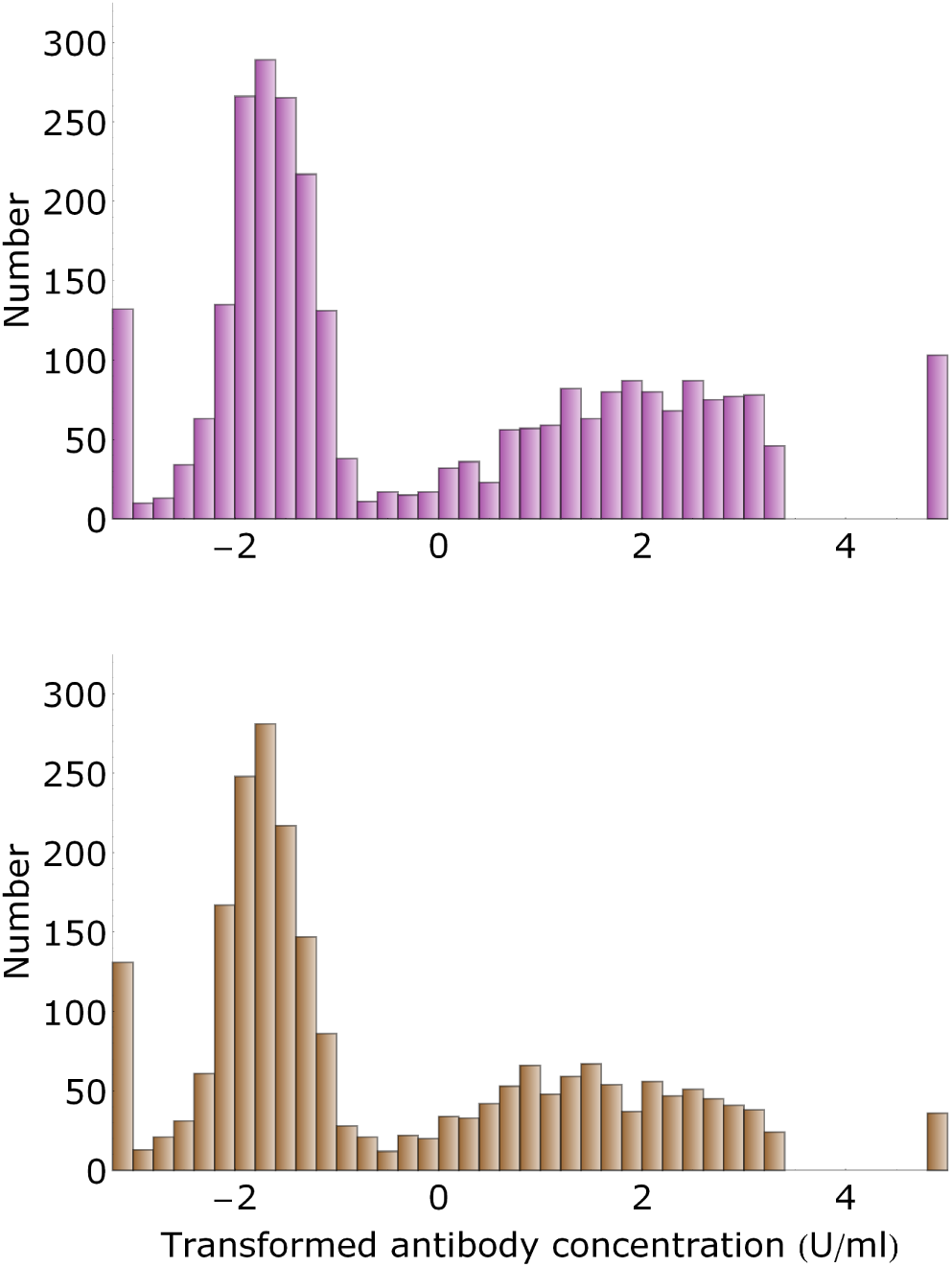
Numbers of samples as a function of antibody concentration. Histograms shows the number of samples by antibody concentration class (top, purple: females; bottom, brown: males). Samples in the bars at 5 U/ml are right-censored at concentration 3.41 U/ml. Total number of female and males samples is 2,842 and 2,337, respectively.

**Fig 8.**
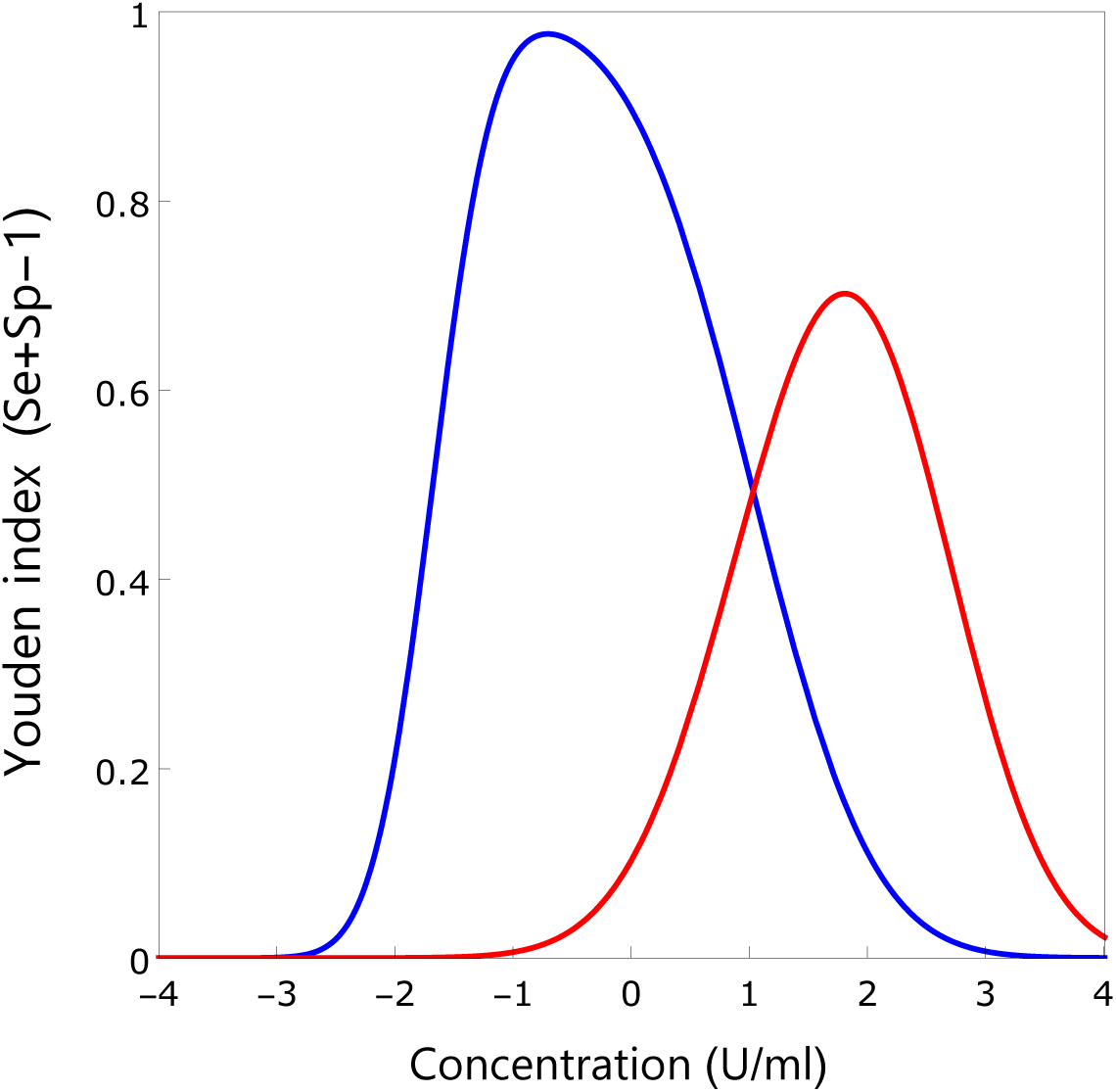
Youden index as a function of antibody concentration. Shown are the Youden indices (Se+Sp-1) for binary classifications, i.e. when using the antibody concentration on the x-axis as cut-off. Classifications are uninfected (S) versus infected (L+B; blue line) and uninfected plus infected with intermediate antibody concentration (S+L) versus infected with increased antibody concentration (B; red line).

**Fig 9.**
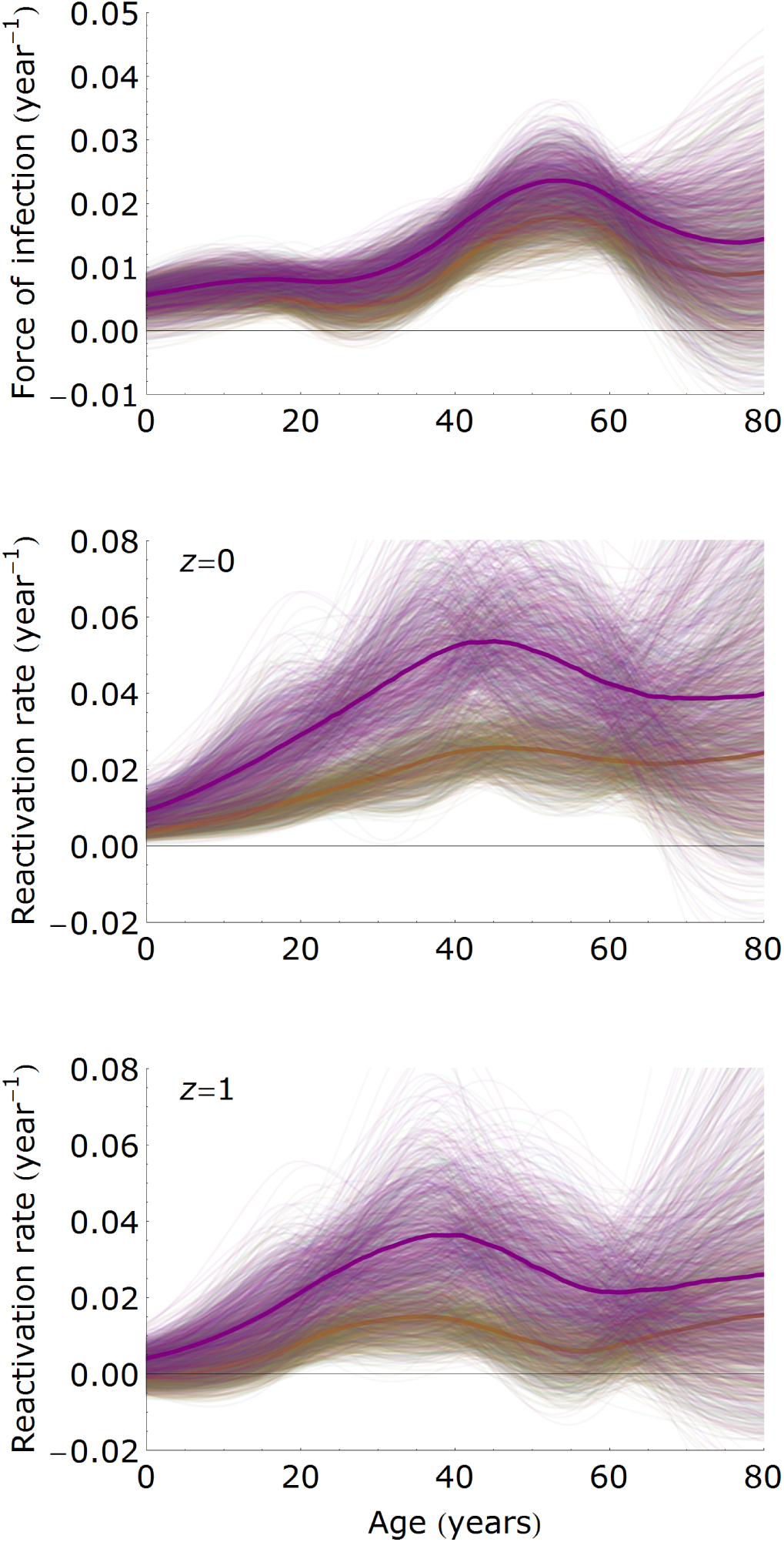
Estimates of transmission parameters with the mixture model. Shown are estimates of the forces of infection (top panel) and reactivation rates (middle and bottom panel) in females (purple) and males (brown) based on spline estimates of the age specific prevalence in the mixture model and Equation (3). The middle panel shows results in the absence of re-infection (*z* = 0), and the bottom panel if re-infection occurs at the same rate as primary infection (*z* = 1). Shown are 1, 000 samples from the posterior distribution (thin lines) with the posterior medians (bold lines). Notice that the force of infection is higher in females than in males, that the reactivation rate is substantially higher in females than in males, and that both the reactivation rates and forces of infection are highest in 30 – 50 year old persons.

**Fig 10.**
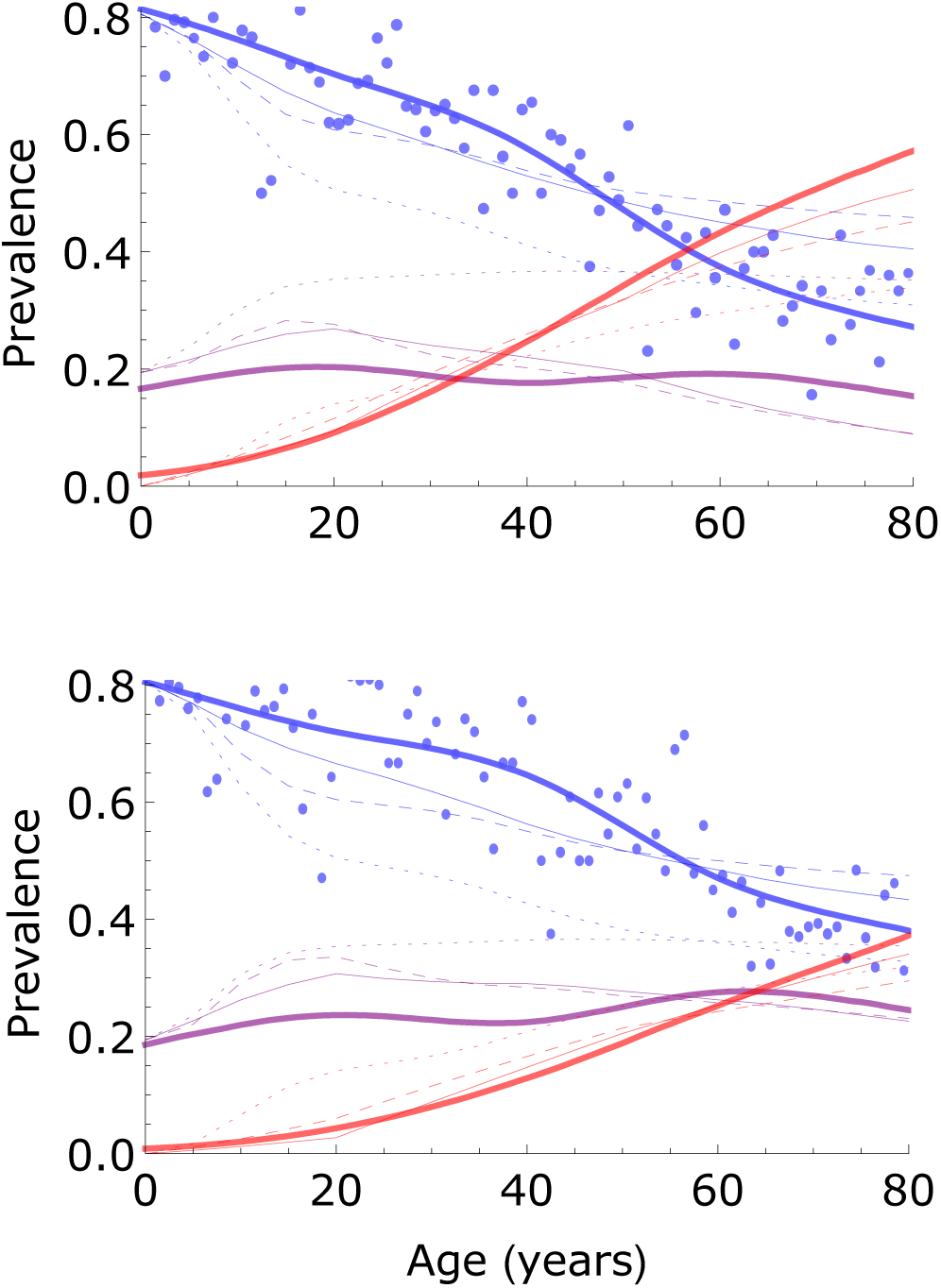
Comparison of fits of the statistical and transmission models. Shown are prevalence estimates for females (top) and males (bottom), and for classes of low (blue), medium (purple), and high (red) antibody measurements. The data and fit (posterior medians) of the statistical model are indicated by dots and thick lines, respectively (cf. Fig 3). Thin solid lines show the fit (using the ML parameter estimates) of the transmission model with infectious reactivation and re-infection. Dotted and dashed lines show the fits of the transmission model with no reactivation and with reactivation and re-infection not being infectious, respectively. The fit of the transmission model with infectious reactivation but no re-infection is very close to the fit of the full model with infectious reactivation and infectious re-infection, and is not shown.

**Fig 11.**
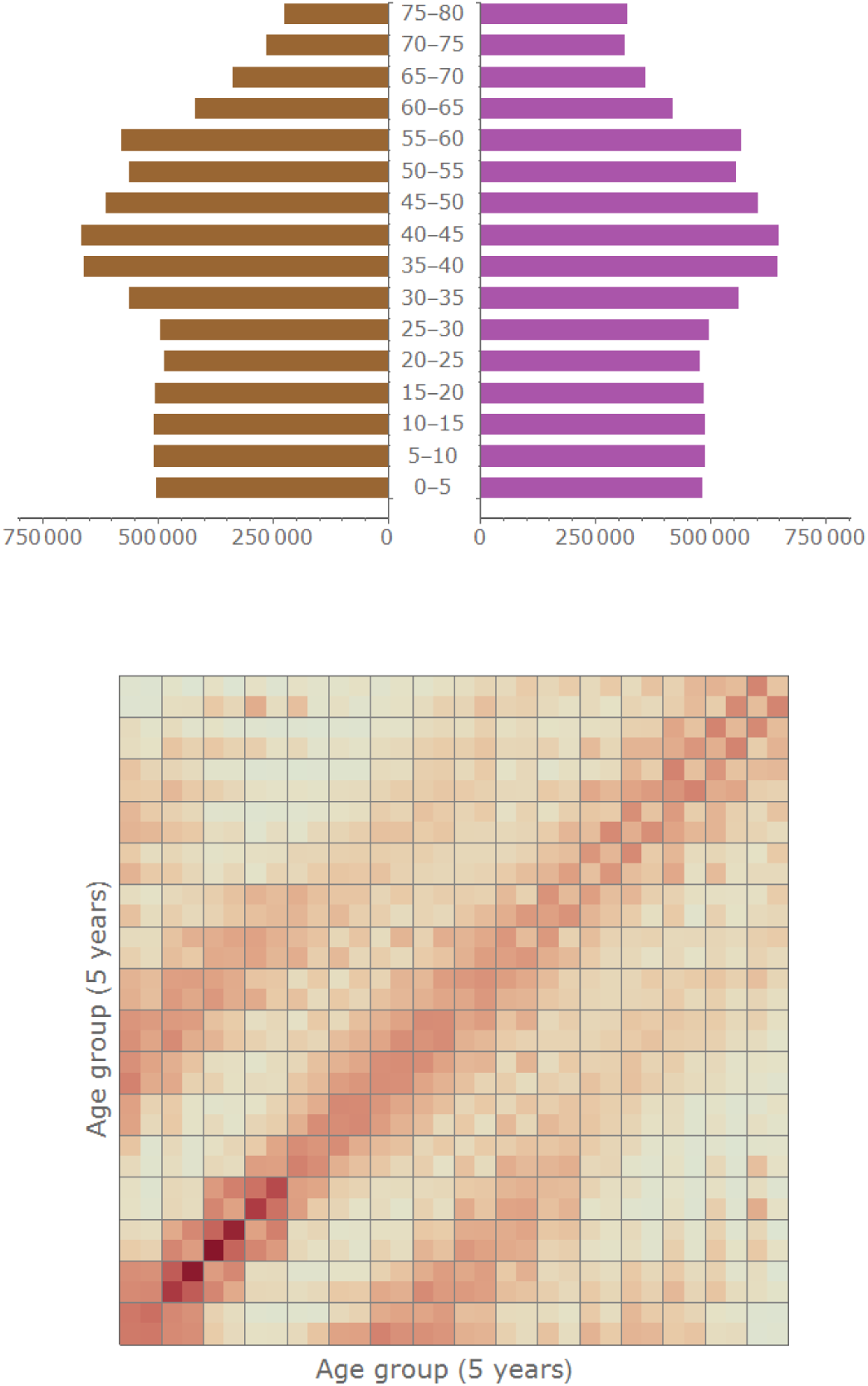
Demography and contact patterns. Shown are the demographic composition of the Dutch population in 2007 and estimated contact rates using data from 2006-2007. The main stratification of contact rates in 5-year age groups, and the secondary stratification by sex (2 by 2 blocks). See [24] for details.

**Fig 12.**
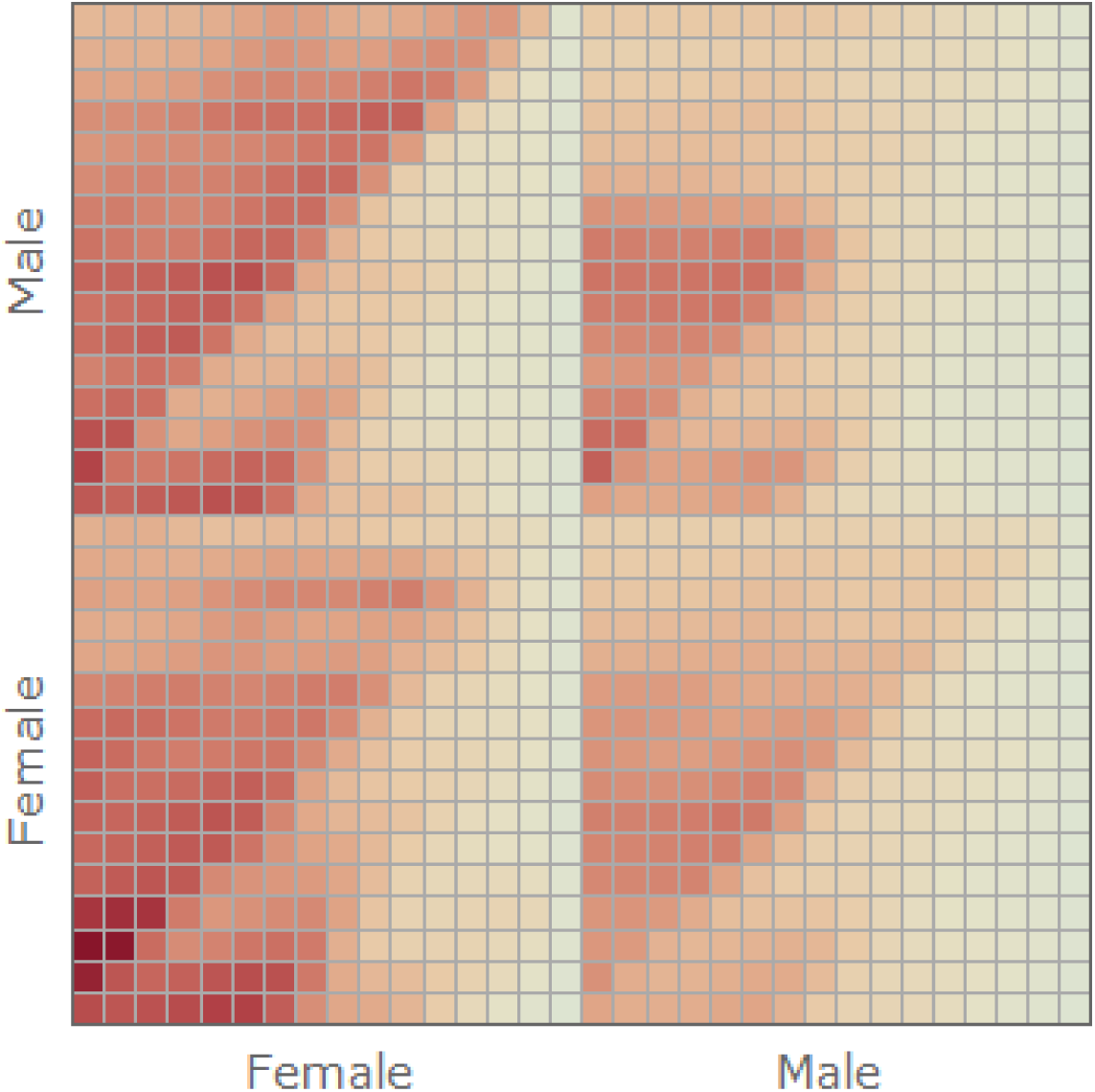
Graphical representation of the next-generation matrix. Shown is the next-generation matrix, stratified by sex and age (16 age groups of 5 years). The top left-hand block shows 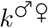, representing the female-to-male transmission 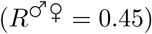. Top right-hand block represents male-to-male transmission 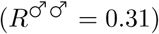. The bottom blocks depict the female-to-female and male-to-female transmission matrices 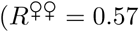 and 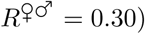. The basic reproduction number is *R*_0_ = 0:83. Orange and blue represent high and low transmission intensity, respectively.

